## Supplemental material for "The *C. elegans* homolog of Sjögren’s Syndrome Nuclear Antigen 1 is required for the structural integrity of the centriole and bipolar mitotic spindle assembly"

Table S1. *C. elegans* strains used in this study

| | |
|---|---|
| N2 | Wild type |
| EU3000 | *sas-7(or1940[gfp::sas-7]) III; ItIs37 [pie-1p::mCherry::his-58 + unc-119(+)] IV* |
| OC210 | *fem-1(hc17) IV* |
| OC779 | *bsSi15 [pKO109: spd-2p-spd-2::mCherry::spd-2 3'-utr, unc-119(+)] I; bsSi30[pCW9: unc-119(+) pcdk-11.2::sfgfp::his-58::cdk-11.2 3' utr] II; unc-119(ed3) III* |
| OC786 | *zyg-1 (bs212 [HA::zyg-1]) II* |
| OC908 | *bsSi30[pCW9: unc-119(+) pcdk-11.2::sfgfp::his-58::cdk-11.2 3' utr] II; bsIs20[pNP99: unc-119(+) tbb-1p::mCherry::tbb-2::tbb-2 3'-utr]; bsIs2 [pCK5.5: Ppie-1::gfp::spd-2]* |
| OC953 | *sas-6 (bs175 [sas-6::HA]) IV* |
| OC955 | *sas-4 (bs177 [HA::sas-4]) III* |
| OC966 | *sas-4 (bs186 [spot::sas-4]) III* |
| OC994 | *sas-4(bs195 [sas-4::gfp]) III* |
| OC1002 | *zyg-1 (bs197[zyg-1::spot]) II* |
| OC1013 | *bsSi30[pCW9: unc-119(+) pcdk-11.2::sfgfp::his-58::cdk-11.2 3' utr] II; bsIs20[pNP99: unc-119(+) tbb-1p::mCherry::tbb-2::tbb-2 3'-utr]; bsIs2 [pCK5.5: Ppie-1::gfp::spd-2]; ssna-1 (bs182) / dpy-9 (tm9713) kvs-5 (tmIs1245)] IV* |
| OC1014 | *ssna-1(bs182) / dpy9 (tm9713) kvs5 (tmIs1245) IV* |
| OC1018 | *ssna-1 (bs206 [ssna-1::spot]) IV* |
| OC1020 | *ssna-1(bs182)/ears-2(ve631[LoxP + myo-2p::gfp::unc-54 3' UTR + rps-27p::neoR::unc-54 3' UTR + LoxP]); fem1 (ht17ts) IV* |
| OC1021 | *zyg-1 (bs197 [zyg-1::spot] II; ssna-1 (bs182) IV* |
| OC1040 | *sas-4(bs195 [sas-4::gfp]) III; ssna-1 (bs206 [ssna-1::spot]) IV* |
| OC1046 | *ssna-1(bs182)/dpy9 (tm9713) kvs5 (tmIs1245) IV* |
| OC1047 | *zyg-1(it25) II* |
| OC1048 | *zyg-1(it25) II; ssna-1(bs182)/dpy9 (tm9713) kvs5 (tmIs1245) IV* |
| OC1052 | *ssna-1 (bs220 [1-99])/dpy9 (tm9713) kvs5 (tmIs1245) IV* |
| OC1068 | *ssna-1 (bs222 [G7X])/dpy9 (tm9713) kvs5 (tmIs1245) IV* |
| OC1070 | *ssna-1 (bs224 [Δ2-22])/dpy9 (tm9713) kvs5 (tmIs1245) IV* |
| OC1072 | *ssna-1 (bs231 [Δ2-17])/dpy9 (tm9713) kvs5 (tmIs1245) IV* |

| | |
|---|---|
| OC1076 | *ssna-1 (bs235 [Δ2-18])/dpy9 (tm9713) kvs5 (tmIs1245) IV* |
| OC1078 | *ssna-1 (bs237[Δ2-18::spot])/dpy9 (tm9713) kvs5 (tmIs1245) IV* |
| OC1126 | *spd-5(vie26[gfp::spd-5 +loxP]) I; ssna-1(bs206 [ssna-1::spot]) IV* |
| OC1129 | *spd-5(vie26[gfp::spd-5 +loxP]) I; ltIs37[pAA64: unc-119(+) pie-1-mcherry-his58]* |
| OC1130 | *spd-5(vie26[gfp::spd-5 +loxP]) I; ssna-1(bs182) IV; ltIs37[pAA64: unc-119(+) pie-1-mcherry-his58]* |
| OC1132 | *ssna-1 (bs243[ssna-1::wrmScarlet) IV* |
| OC1134 | *sas-6 (bs190 [sas-6::spot]) IV* |
| OC1135 | *ssna-1(bs182), sas-6 (bs190 [sas-6::spot]) IV* |
| OC1138 | *ssna-1(bs182)/ears-2(ve631[LoxP + myo-2p::gfp::unc-54 3' UTR + rps-27p::neoR::unc-54 3' UTR + LoxP]) IV* |
| OC1139 | *ssna-1 (bs243 [ssna-1::wrmScarlet]) IV; ltSi560 [pPLG014; Pmex-5::gfp::his-11::tbb2_3'UTR, tbg-1::gfp::tbb-2_3'UTR; cb-unc-119(+)]V* |
| OC1166 | *sas-1(t1476, bs272) III* |
| OC1177 | *zyg-1(it25) II; bsSi15 [pKO109: spd-2p-spd-2::mCherry::spd-2 3'-utr, unc-119(+)] I; bsSi30[pCW9: unc-119(+) pcdk-11.2::sfgfp::his-58::cdk-11.2 3' utr] II* |
| OC1178 | *zyg-1(it25) II; bsSi15 [pKO109: spd-2p-spd-2::mCherry::spd-2 3'-utr, unc-119(+)] I; bsSi30[pCW9: unc-119(+) pcdk-11.2::sfgfp::his-58::cdk-11.2 3' utr] II; ssna-1 (bs182)/ears-2(ve631[LoxP + myo-2p::GFP::unc-54 3' UTR + rps-27p::neoR::unc-54 3' UTR + LoxP]) IV* |
| OC1183 | *bsSi15 [pKO109: spd-2p-spd-2::mCherry::spd-2 3'-utr, unc-119(+)] I; bsSi30[pCW9: unc-119(+) pcdk-11.2::sfgfp::his-58::cdk-11.2 3' utr] II; ssna-1 (bs182)/ears-2(ve631[LoxP + myo-2p::gfp::unc-54 3' UTR + rps-27p::neoR::unc-54 3' UTR + LoxP]) IV* |
| OC1205 | *sas-5(bs199 [sas-5::spot]) V* |
| OC1207 | *ssna-1(bs182)/ears-2(ve631[LoxP + myo-2p::gfp::unc-54 3' UTR + rps-27p::neoR::unc-54 3' UTR + LoxP]) IV; sas-5(bs199 [sas-5::spot]) V* |
| OC1209 | *sas-7(or1940[gfp::sas-7]) III; ssna-1(bs182)/ears-2(ve631[LoxP + myo-2p::gfp::unc-54 3' UTR + rps-27p::neoR::unc-54 3' UTR + LoxP]), ItIs37 [pie-1p::mCherry::his-58 + unc-119(+)] IV* |
| OC1264 | *sas-4 (bs186 [spot::SAS-4]) III; ssna-1(bs182)/ears-2(ve631[LoxP + myo-2p::gfp::unc-54 3' UTR + rps-27p::neoR::unc-54 3' UTR + LoxP]) IV* |
| OC1265 | *sas-4(bs195 [sas-4::gfp]) III; ssna-1(bs182)/ears-2(ve631[LoxP + myo-2p::gfp::unc-54 3' UTR + rps-27p::neoR::unc-54 3' UTR + LoxP]) IV* |

| | |
|---|---|
| OC1335 | *sas-1(t1476, bs272) III; ssna-1(bs182)/ears-2(ve631[LoxP + myo-2p::gfp::unc-54 3' UTR + rps-27p::neoR::unc-54 3' UTR + LoxP]) IV* |
| OC1344 | *ssna-1 (bs356 [R18E::spot]) IV* |

Table S2: crRNAs and repair templates used in this study

| Allele | Background | crRNA (5' -> 3') | Repair template (5' -> 3') |
|---|---|---|---|
| bs199 | N2 | TTCGTGAAAAATACGCTCGC | CTGAACGAGAACGCCGTATTCGTGAAAAATACGCTCGC<br>AGAAAACCAGACCGTGTCCGTGCCGTCTCCCACTGGT<br>CCTCCTGATATCAAATGTGTTTAACTCTTGACGTTTTAAA |
| bs206 | N2 | CTTTGTGCGCAAAGAGTATC<br>GATATATTTACAGGCATTTT | GCAAAAGACGTTGGTGGACTTTGTGCGCAAAGAGTATC<br>AAGATACGAAACATCAGAAATATCCAGACCGTGTCCGTG<br>CCGTCTCCCACTGGTCCTCCTGAACTCTGAACAACTGTC<br>TCCCAAAAATGCCTGTAAATATATCAATTATCGACATAACTTC |
| bs233 | bs231 | | |
| bs237 | bs235 | | |
| bs249 | bs217 | | |
| bs357 | bs286 | | |
| bs218 | N2 | | GCAAAAGACGTTGGTGGACTTTGTGCGCAAAGAGTATCAA<br>GATACGAAACATCAGAAATATGAACCGGAAGCGTGAACTCT<br>GAACAACTGTCTCCCAAAAATGCCTGTAAATATATCAATTATC<br>GACATAACTTC |
| bs242 | N2 | | GCAAAAGACGTTGGTGGACTTTGTGCGCAAAGAGTATCAA<br>GATACGAAACATCAGAAATATGGATCCGCCGGATCCGCCG<br>CCGGATCCGGAGAGTTCGTCAGCAAGGGAGAGGCAGTTAT<br>CAAGGAGTTCATGCGTTTCAAGGTCCACATGGA |
| | | | TATCAAGGAGTTCATGCGTTTCAAGGTCCACATGGAGGGAT<br>CCATGACCGAGGGACGTCACTCCACCGGAGGAATGGACGA<br>GCTCTACAAGTGAACTCTGAACAACTGTCTCCCAAAAATGCC<br>TGTAAATATATCAATTATCGACATAACTTC |
| bs243 | bs242 | CATGGAGGGATCCATGACCG | GTCAGCAAGGGAGAGGCAGTTATCAAGGAGTTCATGCGTTT<br>CAAGGTCCACATGGAGGGATCCATGAACGGACACGAGTTCG<br>AGATCGAGGGAGAGGGAGAGGGACGTCCATACGAGGGAAC<br>CCAAACCGCCAAGCTCAAGGTCACCAAGGGAGGACCACTC<br>CCATTCTCCTGGGACATCCTCTCCCCACAATTCATGTACGGA<br>TCCCGTGCCTTCACCAAGCACCCAGCCGACATCCCAGACTA<br>CTACAAGCAATCCTTCCCAGAGGGATTCAAGTGGGAGCGTG<br>TCATGAACTTCGAGGACGGAGGAGCCGTCACCGTCACCCAA<br>GACACCTCCCTCGAGGACGGAACCCTCATCTACAAGGTCAA<br>GCTCCGTGGAACCAACTTCCCACCAGACGGACCAGTCATGC<br>AAAAGAAGACCATGGGATGGGAGGCCTCCACCGAGCGTCTC<br>TACCCAGAGGACGGAGTCCTCAAGGGAGACATCAAGATGGC<br>CCTCCGTCTCAAGGACGGAGGACGTTACCTCGCCGACTTCA<br>AGACCACCTACAAGGCCAAGAAGCCAGTCCAAATGCCAGGA<br>GCCTACAACGTCGACCGTAAGCTCGACATCACCTCCCACAAC<br>GAGGACTACACCGTCGTCGAGCAATACGAGCGTTCCGAGGG<br>ACGTCACTCCACCGGAGGAATGGACGAGCTCTACAAG |
| bs182 | N2 | TAGAATCATGCATTTGCATT<br>CTTTGTGCGCAAAGAGTATC | TTCGTATTTGAACAATTACTGACTAATTTCCTCCGAATGCAAA<br>TGCATGATTCTAGAACAAAAAAAACATCAGAAATATTGAACTC<br>TGAACAACTGTCTC |

| | | | |
|---|---|---|---|
| bs215 | N2 | CTGTGAGACGGCGTTCCTCT | GAAGATAAACATTTATAGTAATATTTCAGACATCCAAGCGCTCAGAGAGGAACGCCGTCTCACAGAATCGTCGATTCGAAAAATG |
| bs216 | bs215 | AGACATCCAAGCGCTCGCGG | GTAAGAGCAAATTGAAGATAAACATTTATAGTAATATTTCAGACATCCAAGCGCTCGCGGAAGAACGCCGTCTCACAGAATCGTCGATTCGAAAAATGGA |
| bs217 | | | GTAAGAGCAAATTGAAGATAAACATTTATAGTAATATTTCAGACATCCAAGCGCTCGCGGCAGCACGCCGTCTCACAGAATCGTCGATTCGAAAAATGGAAA |
| bs220 | N2 | CTTTGTGCGCAAAGAGTATC | GCAAAAGACGTTGGTGGACTTTGTGCGCAAAGAGTATCAAGATTGAACTCTGAACAACTGTCTCCCAAAAATGCCTGT |
| bs222 | N2 | AAAATGTCTTCTCGATCTAC | TGCATGATTCTAGAACAAAAAATGTCTTCTCGATCTACATGAAGCTTTGATGAAATATCACAGTGTAAGAGCAAATT |
| bs224 | bs215 | AAATGTCTTCTCGATCTAC<br>AGACATCCAACGTCTCCGAG | CCGAATGCAAATGCATGATTCTAGAACAAAAAATGCGCCGTCTCACAGAATCGTCGATTCGAAAAATGGA |
| bs231 | N2 | AAAATGTCTTCTCGATCTAC<br>CTGTGAGACGGCGTTCCTCT | CCGAATGCAAATGCATGATTCTAGAACAAAAAATGCGTCTCAGAGAGGAACGCCGTCTCACAGAATCGTCGATTCGAAAAA |
| bs235 | N2 | | CCGAATGCAAATGCATGATTCTAGAACAAAAAATGCTCAGAGAGGAACGCCGTCTCACAGAATCGTCGATTCGAAAAA |
| bs284 | N2 | | TGCATGATTCTAGAACAAAAAATGTCTTCTCGAAGCACAGGAAGCTTTGATGAAATATCACAGGGTAAGAGCAAATTGAAGATAAACATTTATAGTAATATTTCAGAGATCCAACGTCTCAGAGAGGAACGCCGTCTCACAGAATCGTCGATTCGAAAAA |
| bs286 | N2 | CTGTGAGACGGCGTTCCTCT | GAAGATAAACATTTATAGTAATATTTCAGACATCCAAGAGCTCAGAGAGGAACGCCGTCTCACAGAATCGTCGATTCGAAAAATG |
| bs312 | bs284 | CTTTGTGCGCAAAGAGTATC | CGCAAAAGACGTTGGTGGACTTTGTGCGCAAAGAGGAGCAGGATACGAAACATCAGAAATATTGAACTCTGAACAAC |
| bs314 | bs286 | | |
| bs335 | N2 | | |

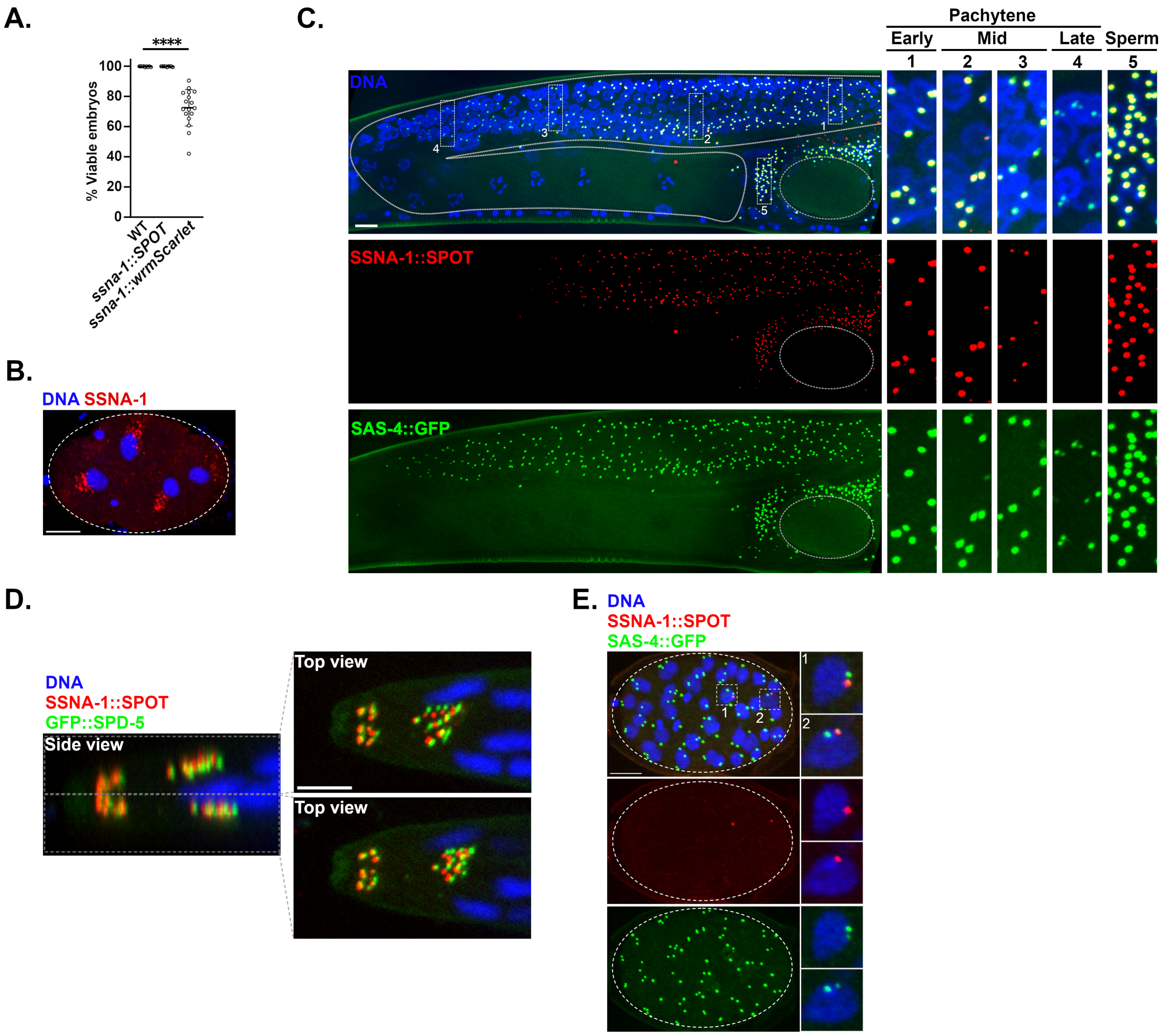


**Fig. S1: SSNA-1 is a stable component of centrioles with broad distribution in the worm. A.**

Quantification of embryonic viability of strains expressing different C-terminal epitope tagged versions of endogenous SSNA-1. Each datapoint represent progeny of a single hermaphrodite. **B.** Representative image of an embryo stained for endogenous SSNA-1 and DNA. Bar = 10 µm. **C.** Localization of SSNA-1::SPOT in the germline with SAS-4::GFP marking centrioles, showing that SSNA-1 is present on centrioles on immature female germ cells as well as sperm. SSNA-1 is lost from germ cell centrioles during mid-pachytene (box 3), prior to SAS-4 (box 4). Bar = 10 µm. **D.** SSNA-1::SPOT is found adjacent to the acentriolar centrosome (marked with GFP::SPD-5) in amphid neurons of L1 larvae. Bar = 10 µm. **E**. An embryo produced by mating SSNA-1::SPOT males to SAS-4::GFP hermaphrodites stained for SAS-4::GFP (green), SSNA-1::SPOT (red) and DNA (blue). Even after many cell cycles SSNA-1::SPOT remains associated with the two original sperm centrioles revealing that SSNA-1 is a stable component of centrioles. Scale bar = 10 µm.

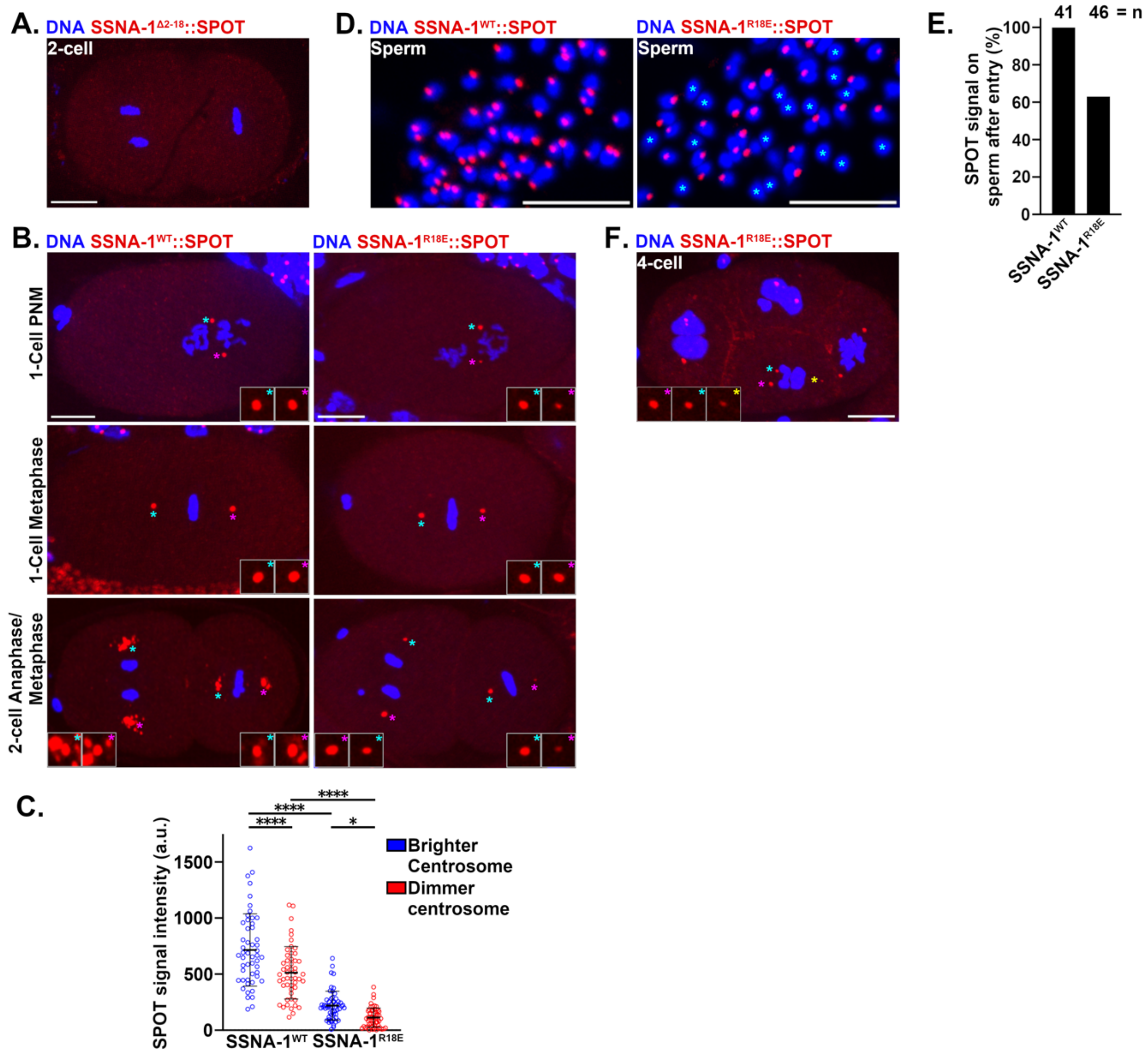


**Fig. S2: SSNA-1 oligomerization and MT binding are required for centriolar and satellite localization. A.** Representative image of a two-cell stage embryo expressing SSNA-1$^{\Delta 2\text{-}18}$::SPOT showing that SSNA-1 fails to localize to the centriole and satellite-like structures. Scale bar = 10 μm. **B.** Localization of SSNA-1$^{WT}$::SPOT (left column) and SSNA-1$^{R18E}$::SPOT (right column)

from pronuclear migration (PNM) to the late 2-cell stage in strains grown at 20°C. Full-size images are maximum intensity projections while insets are 2X magnifications of the center z plane of each centrosome indicated by colored asterisks. Scale bar = 10 μm. **C.** Quantification of SSNA-1 signal intensity from B showing that the centriole levels of SSNA-1$^{R18E}$::SPOT are strongly reduced relative to wild-type. Also notice that in both the wild-type and mutant, that one centrosome is brighter than the other. **D.** Maximum intensity projections of sperm from SSNA-1$^{WT}$::SPOT and SSNA-1$^{R18E}$::SPOT strains. Note that in the wild type, all sperm nuclei are associated with a SSNA-1::SPOT focus while many sperm in the SSNA-1$^{R18E}$::SPOT strain lack a focus (asterisks). Scale bar = 10 μm. **E.** Quantitation showing the percentage of wild-type and R18E mutant sperm that possess a focus of SSNA-1. **F.** A four-cell stage embryo expressing SSNA-1$^{R18E}$::SPOT grown at 25°C. Note the presence of a multipolar spindle in the EMS cell at bottom. Full-scale images are maximum intensity projections while insets are 2X magnifications of the center z plane of each centrosome as marked by colored asterisks. Scale bar = 10μm.

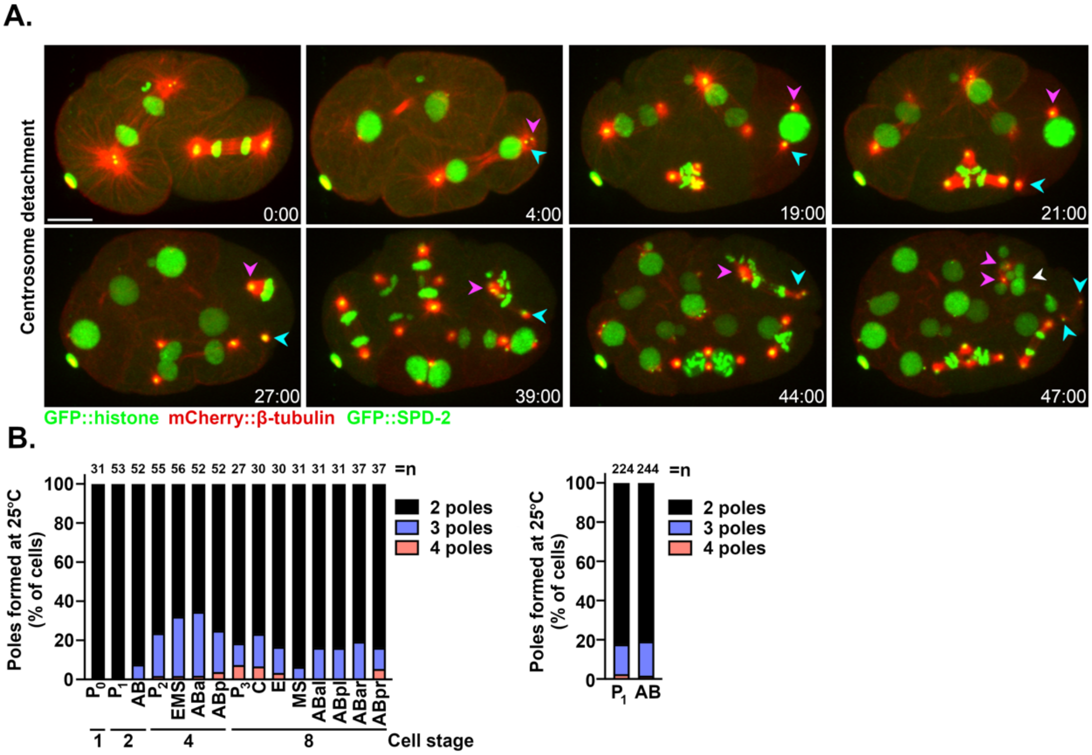


**Fig. S3: Cell division defects during development of *ssna-1(Δ)* embryos. A.** Frames from a time-lapse recording of an *ssna-1(Δ)* embryo expressing GFP::SPD-2 (green); GFP::histone (green); and mCherry::β-tubulin (red). Magenta and cyan arrowheads indicate the two centrosomes that arise from one pole of the $P_1$ spindle (t=4:00). One centrosome stays associated with the $P_2$ nucleus (magenta) while the other (cyan) migrates to the cell periphery (t=21:00). The itinerant centrosome makes a late contribution to bipolar spindle assembly (t=44:00), but chromosome segregation defects lead to micronuclei formation (white arrowhead) (t=47:00). Note that both centrosomes duplicated. Scale bar = 10 μm. **B.** Distribution of multipolar spindle formation among the early embryonic blastomeres (n= number of cells scored). **C.** Percentage of cells within the $P_1$ and AB lineages that form multipolar spindles.

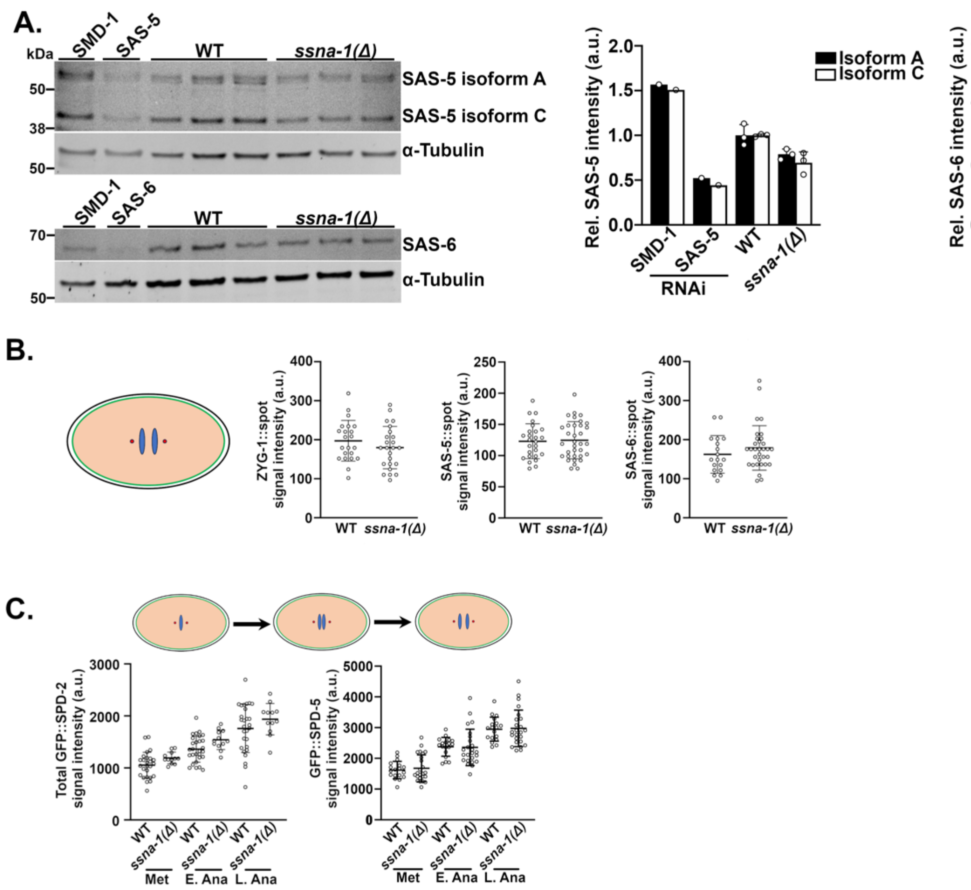


**Fig. S4. Loss of SSNA-1 is not associated with overexpression of centriole duplication factors. A.** Immunoblots of SAS-5 and SAS-6 levels in extracts prepared from gravid hermaphrodites (left). Quantitation of blots (right) shows the relative levels after normalization to α-tubulin. **B.** Centriole-associated levels of ZYG-1::SPOT, SAS-5::SPOT, and SAS-6::SPOT at anaphase in the zygote. **C**. Centrosome-associated levels of GFP::SPD-2 (left) and GFP::SPD-5 (right) from metaphase through anaphase in the zygote, as measured by quantitative immunofluorescence. In panels B and C, all differences between wild-type and *ssna-1(Δ)* mutants are nonsignificant.

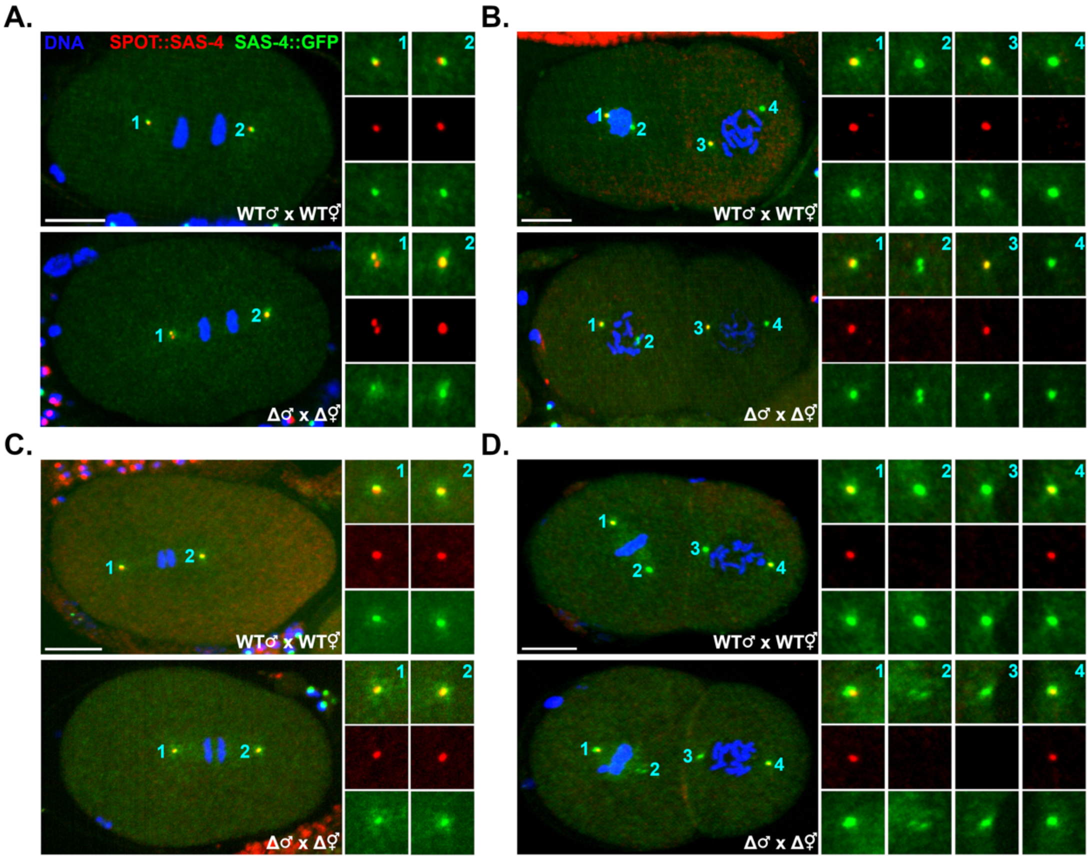


**Fig S5. Loss of *ssna-1* results in centriole fragmentation rather than premature disengagement. A-D.** Representative images of embryos produced from *SPOT::sas-4* male x *sas-4::GFP* hermaphrodite matings. Insets are 2X magnifications. All images are maximum intensity projections. Scale bars = 10 µm**. A.** WT and *ssna-1(Δ)* embryos at mid anaphase. In WT embryos, mother (red) and daughter (green) centrioles remain associated prior to normal disengagement in late anaphase/telophase while in *ssna-1(Δ)* embryos, mother (red) centrioles fracture (centrosome 1). One of these two red centriole fragments remains associated with the green daughter centriole indicating the centriole pair is still engaged. **B.** WT and *ssna-1(Δ)* embryos during prophase and prometaphase. Note that all red centrioles and their green daughters are coincident and therefore engaged. Note also an instance of apparent centriole fragmentation of a green centriole in the *ssna-1(Δ)* embryo (centrosome 2). **C and D.** Representative images of one- (C) and two-cell (D) embryos showing that in both wild-type and *ssna-1(Δ)* embryos, all red paternally derived centrioles remain engaged with their gene daughters through early anaphase (WT = 64 centrosomes and *ssna-1(Δ)* = 56 centrosomes).

**Movie S1. SSNA-1 satellite-like structures display dynamic behavior.** Time-lapse video corresponding to Fig. 2E of an embryo expressing SSNA-1::wrmScarlet, GFP::histone, and γ-tubulin::GFP. A side view of the ABpl blastomere dividing is shown. SSNA-1 satellites display dynamic behavior during division whereby they disperse at the end of mitosis following PCM breakdown and spread out across the nucleus. As the cell progresses through mitosis, the satellites accumulate around the centrosome of each pole. Maximum accumulation is achieved by late metaphase/early anaphase.
